# Identification of a microbial sub-community from the feral chicken gut that reduces *Salmonella* colonization and improves gut health in a gnotobiotic chicken model

**DOI:** 10.1101/2022.07.23.501241

**Authors:** Supapit Wongkuna, Roshan Kumar, Sudeep Ghimire, Samara Mattiello Drescher, Abhijit Maji, Achuthan Ambat, Linto Antony, Surang Chankhamhaengdecha, Tavan Janvilisri, Eric Nelson, Kinchel C. Doerner, Melissa Behr, Joy Scaria

## Abstract

A complex microbial community in the gut generally prevent the colonization of enteric pathogens such as *Salmonella*. Because of the high complexity, several species or combination of species in the gut can confer colonization resistance. To gain a better understanding of the colonization resistance against *Salmonella enterica*, we isolated a library of 1,300 bacterial strains from feral chicken gut microbiota which represented a total of 51 species. Using a co-culture assay, we screened the representative species from this library and identified 30 species that inhibited *Salmonella enterica* Typhimurium. To improve the *Salmonella* inhibition capacity, from a pool of fast-growing species, we formulated 66 bacterial blends, each of which composed of 10 species. Bacterial blends were more efficient in inhibiting *Salmonella* as compared to individual species. The blend that showed maximum inhibition (Mix10) also inhibited other serotypes of *Salmonella* frequently found in poultry. The *in vivo* effect of Mix10 was examined in a gnotobiotic and conventional chicken model. The Mix10 consortium reduced *Salmonella* colonization, intestinal tissue damage and inflammation in both models. Cell free supernatant of Mix10 did not show *Salmonella* inhibition, indicating that Mix10 inhibits *Salmonella* through either nutritional competition or reinforcement of host immunity. Out of ten species, three species in Mix10 did not colonize while three species constituted more than 70% of the community. Two of these species represents previously uncultured bacteria. Our approach could be used as a high-throughput screening system to identify additional bacterial sub-communities that confer colonization resistance against enteric pathogens and its effect on the host.

**Importance:** *Salmonella* colonization in chicken and human infections originating from *Salmonella-* contaminated poultry is a significant problem. Poultry has been identified as the most common food linked to enteric pathogen outbreaks in the United States. Since multi-drug resistant *Salmonella* often colonize chicken and cause human infections, methods to control *Salmonella* colonization in poultry are needed. The method we describe here could form the basis of developing gut microbiota-derived bacterial blends as a microbial ecosystem therapeutic against *Salmonella*.

## Introduction

A dense and complex microbial community colonizes the human and animal gastrointestinal tract over time. This complex community collectively called the gut microbiota provides a range of functions such as the development of the immune system, digestion, tissue integrity, vitamin and nutrient production, and the ability to prevent colonization of enteric pathogens [1-3]. With the advances in the microbiome research and because of the worldwide increase in the bacterial antibiotic resistance [4], there is high interest in using mature gut microbiome as an alternative means of suppressing enteric infections [5-7]. The ability of the healthy gut microbiota to prevent pathogen colonization was first demonstrated by Nurmi and Rantala in a classic experiment in which inoculation of young chicken with adult chicken feces prevented the colonization of *Salmonella* [8, 9]. The same concept was used in recent years to treat recurrent *Clostridum difficile* infection in humans by fecal transplantation from healthy individuals [10, 11]. Recently, rather than using the whole fecal microbial community, there were efforts to identify individual species in the microbiome that confer colonization resistance [12, 13]. This approach of using single species or combinations of species conferring colonization resistance to prevent pathogen colonization and infection is termed as precision microbiome reconstitution [14] or microbial ecosystem theraputics [15].

Although colonization resistance of the gut microbiota was first demonstrated, *Salmonella* colonization in poultry continues to be a significant problem even today. Poultry has been identified as the most common food in outbreaks with pathogens in the United States [16, 17]. The poultry industry has responded to this problem by implementing biosecurity measures that are designed to minimize exposure of chickens to the pathogens [18]. Conversely, increased biosecurity and clean conditions in the production system would also decrease the exposure to commensal bacteria and would reduce the microbiome diversity in the chicken gut. As proposed by Rolf Freter in 1983 in his nutrient niche hypotheis [19], reduced exposure to commensal gut microbes would open nutrient niches in the gut that can be easily used by pathogens which increases their colonization risk. To reduce this risk, poultry industry has attempted to reduce the pathogen colonization by inoculating chicken with complex commensal bacterial blends such as the lyophilized mixture of anaerobic bacteria from the cecum of adult chicken [20], collection of more than 200 bacteria from pathogen free-birds [21], bacteria from healthy chicken mucosal scrapping [22], and continuous flow culture of cecal chicken bacteria [23]. Due to the complexity of these mixtures, it is difficult to understand their mechanism of action and improve their efficacy.

Recently, a modified Koch postulate was proposed for the mechanistic understanding of the gut microbiota colonization resistance [24, 25]. As per this proposal, a mechanistic study of the colonization resistance require the development of commensals as a pure culture library and then mono or poly-associate them in a germfree host to demonstrate the amelioration of disease. In this study, we used this framework to better understand the colonization resistance of chicken gut microbiota. To this end, we developed a commensal bacterial culture library from *Salmonella*-free feral chicken and formulated a defined bacterial sub-community from *Salmonella* inhibiting commensal species. This consortium was tested to demonstrate the *Salmonella* exclusion capacity using a gnotobiotic chicken model. Our results showed that the defined consortium reduced *Salmonella* colonization, the severity of intestinal tissue damage and inflammation. With further improvements, the current approach and the blend we demonstrate here could offer a means of formulating defined communities of commensal bacteria as microbial ecosystem therapeutic in poultry.

## Results

### Development of the feral chicken gut bacterial library

It is known that microbiota from healthy adult chicken could inhibit the growth of *S. enterica* in the gut [8]. Because of the exposure to a broad range of environmental conditions, feral chicken would have more diverse gut microbiome than commercial chicken and a high percentage of the microbiota in the feral chicken gut could have inhibitory capacity against *S. enterica*. To ascertain this, we isolated a bacterial library from the pooled intestinal contents of six feral chicken using anaerobic culture conditions. We used a modified Brain Heart Infusion as the base culture medium which is hereafter referred as BHI-M (Table S1). When a non-selective medium is used for cultivation, it is common that fast-growing bacteria use up space and nutrients in the medium. To avoid this problem, we used iterative antibiotic supplementation of BHI-M to suppress bacteria that dominated the base medium (Table S1). For example, from the base BHI-M, when 32 bacterial species were isolated, five species (*Massiliomicrobiota timonensis, Faecalicoccus pleomorphus, Eubacterium cylindroides, Collinsella* sp., and *Olsonella* sp.) accounted for 52.6% of picked colonies. To suppress the growth of these species, we supplemented BHI-M with gentamycin and kanamycin which allowed isolation of several species that was not isolated from the plain medium. Using 12 such selection conditions, 1,300 isolates were selected. Species identity of 1,023 isolates was determined by either MALDI-TOF or 16S rRNA gene sequencing (Table S2). Fig. 1 shows an overview of the culture conditions, diversity, and frequency of the isolated species. Overall, we identified 51 species using a cut off of 97.82% at 16S rRNA gene sequence level. The identified species were mostly dominated by the phylum *Firmicutes* (36 species), followed by *Bacteroides* (5 species), *Proteobacteria* (5 species), *Actinobacteria* (4 species) and *Fusobacterium* (1 species). Altogether, we also captured 11 previously uncultured species which can represent the novel type strains.

**Figure 1.**
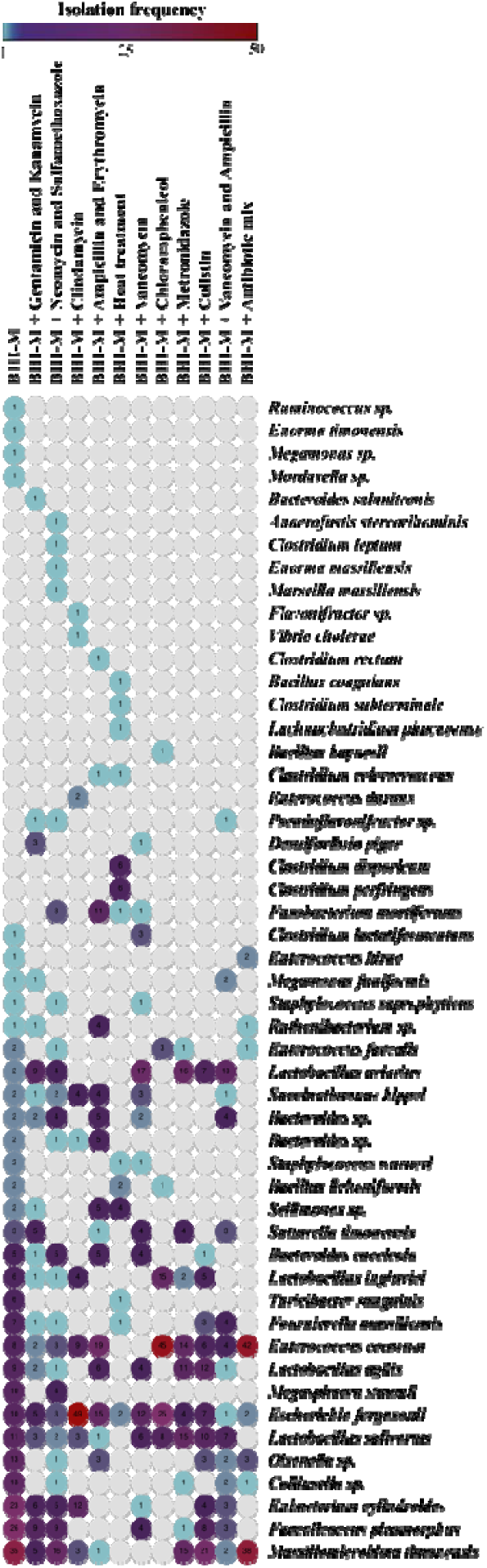
Diversity and frequency of bacterial species isolated from the feral chicken gut microbiota in culture library. The abundance and diversity of 51 bacterial species (1,023 isolates) were accounted according to culture conditions. Intestinal content of six feral chickens was pooled, stocked, and cultured using 12 culture combinations given in Supplemental Table 1. Species identification was performed using matrix-assisted laser desorption/ionization-time of flight (MALDI-TOF) or 16S rRNA sequencing. A heat map showed diversity and abundance of bacterial species in a culture library was generated using Morpheus, versatile matrix visualization and analysis software. The numbers in each circle represent the frequency of isolation of that species. Full list of strains is given in Supplemental Table 2.

### Screening and the selection of defined bacterial consortium that inhibits Salmonella

To determine the species that could inhibit *Salmonella* in our culture library, we tested the inhibition capacity of representative isolates of 51 species against *S. enterica* Typhimurium (*S*. Typhimurium) using a co-culture assay. From the total collection, 30 species showed varying degree of inhibition against *S*. Typhimurium (Fig. 2A). Since the reduction in pH during bacterial growth is inhibitory to *S*. Typhimurium, we also determined whether pH was reduced at the end of the co-culture assay (Table S3). The pH range varied between 5.5 and 7.0 and in the majority of the cases, pH did not drop below 6.0. This can be explained that the inhibition of *S*. Typhimurium by these strains may not be primarily mediated by the production of organic acids that would have lowered the pH of the medium. Interestingly, this screen also showed that 11 species in our collection enhanced the growth of *S*. Typhimurium (Fig. 2A). Further, we tested whether the *Salmonella* inhibition capacity of these strains have improved when a subset of strains is pooled together. To reduce the complexity of the pool, twelve inhibitory bacterial strains that are fast growing and maintaining a pH above 5.8, were selected to formulate bacterial blends (Table S4). Since there is the chance that species composition of the blend may positively or negatively influence the *S*. Typhimurium inhibitory ability, we made several subsets using a combinatorial approach, in which two species were randomly removed from the 12 selected species. With this combinatorial approach, total 66 different blends, each of which composed of 10 species could be generated (Table S4). We then tested the *S*. Typhimurium inhibitory ability of all these blends using co-culture assay. As shown in Fig. 2B, the blend approach improved the *S*. Typhimurium inhibition. Out of 66 blends, blend 63 showed the highest inhibition with about 250-fold reduction of *S*. Typhimurium compared to control. The majority of blends were inhibitory in varying degrees, while blend 32 and 59 increased the growth of *S*. Typhimurium. This was unexpected because all the selected strains were individually inhibiting *Salmonella*. It is an indication that the community composition of the bacterial blends can override the individual strain phenotype (*Salmonella* inhibition in this case). Therefore, the combinatorial testing we performed can reveal gut microbiota sub-community that might produce an entirely different phenotype than that of the individual species membership in a bacterial consortium. Since blend 63 showed the highest inhibition of *Salmonella* among all tested blends, we further verified properties of this blend *in vivo* experiments. This blend which hereafter referred to as Mix10 (Table 1) was composed of *Faecalicoccus pleomorphus, Lactobacillus agilis, Staphylococcus saprophyticus, Bacillus paralicheniformis, Enterococcus durans, Olsenella* sp., *Megasphaera statonii, Pseudoflavonifractor* sp., and *Massiliomicrobiota timonensis*. Based on 16S rRNA gene similarity search against EzTaxon and NCBI databases [26], two strains (*Olsenella* sp. and *Pseudoflavonifractor* sp.) in this blend represented uncultured organisms of their respective genera of which, a novel species *Olsenella lakotia* SW165^T^ was characterized previously [27]. These results are consistent with our reasoning that feral chicken gut harbor diversity that includes new taxa that are inhibitory against *Salmonella*. To determine whether Mix10 could inhibit other serotypes of *Salmonella* that commonly colonize chicken, we tested the inhibition of Mix10 against *Salmonella* Heidelberg, *Salmonella* Infantis, *Salmonella* Enteritidis and *Salmonella* Typhimurium using the same co-culture assay. These assays revealed that Mix10 inhibits tested *Salmonella* serotypes at similar levels (Fig. S1). To determine the mechanism of Mix10 inhibition, cell free supernatants were tested against *Salmonella* after heat and proteinase K treatment. Nevertheless, our results show that inhibitory activity is lost in the cell free supernatant indicating that inhibition is mediated through nutrient competition (Fig S2).

**Table 1.**
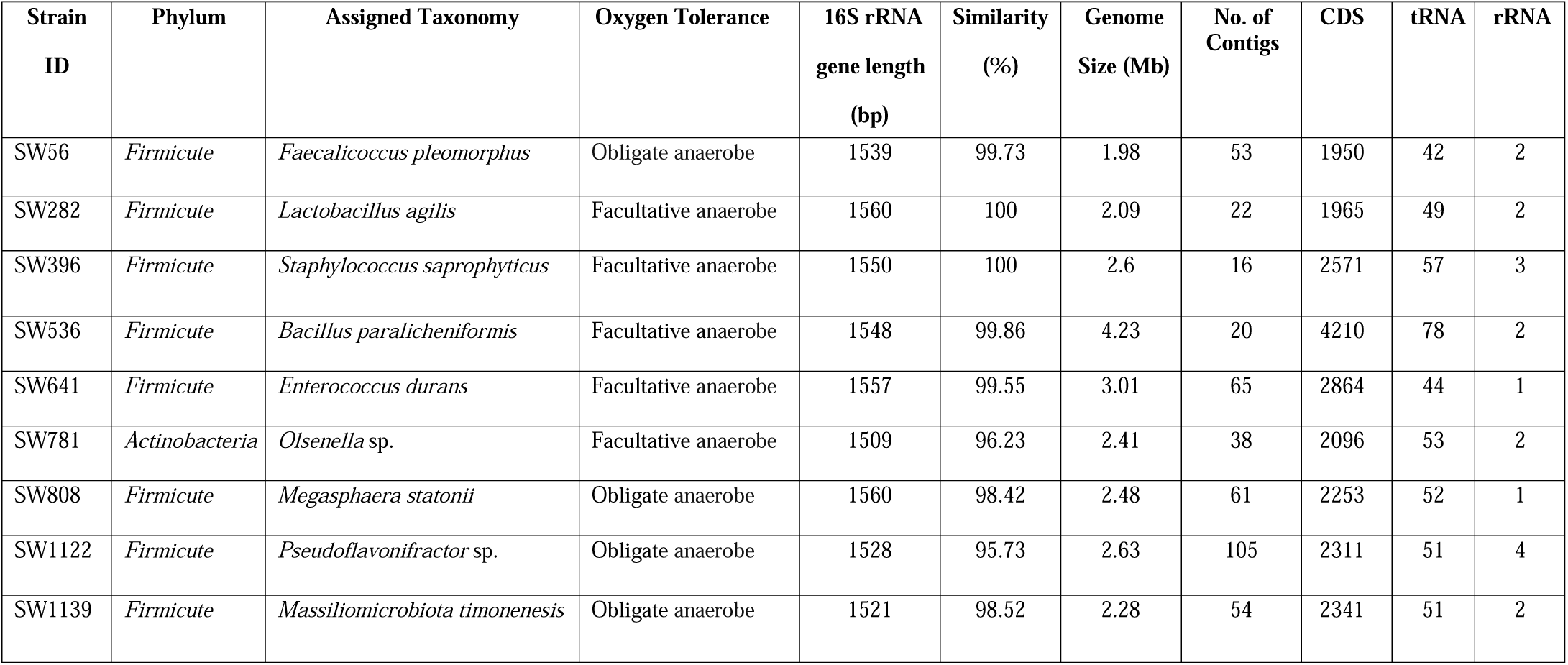
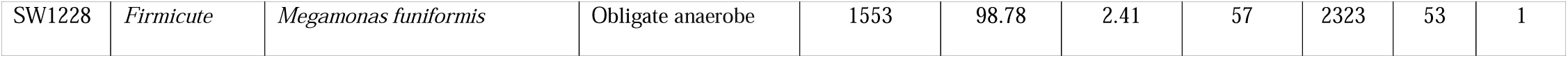
Description of bacterial strains used to formulate Mix10. Whole genome sequencing of each species was performed using Illumina MiSeq. Genome assembly was performed using Unicycle that builds an initial assembly graph from short reads using the *de novo* assembler SPAdes 3.11.1. The open reading frames (ORFs) were predicted using Prodigal 2.6 implemented in the Prokka software package. Species identity of the individual species was determined by searching against EzTaxon using full-length 16S rRNA sequence.

**Figure 2.**
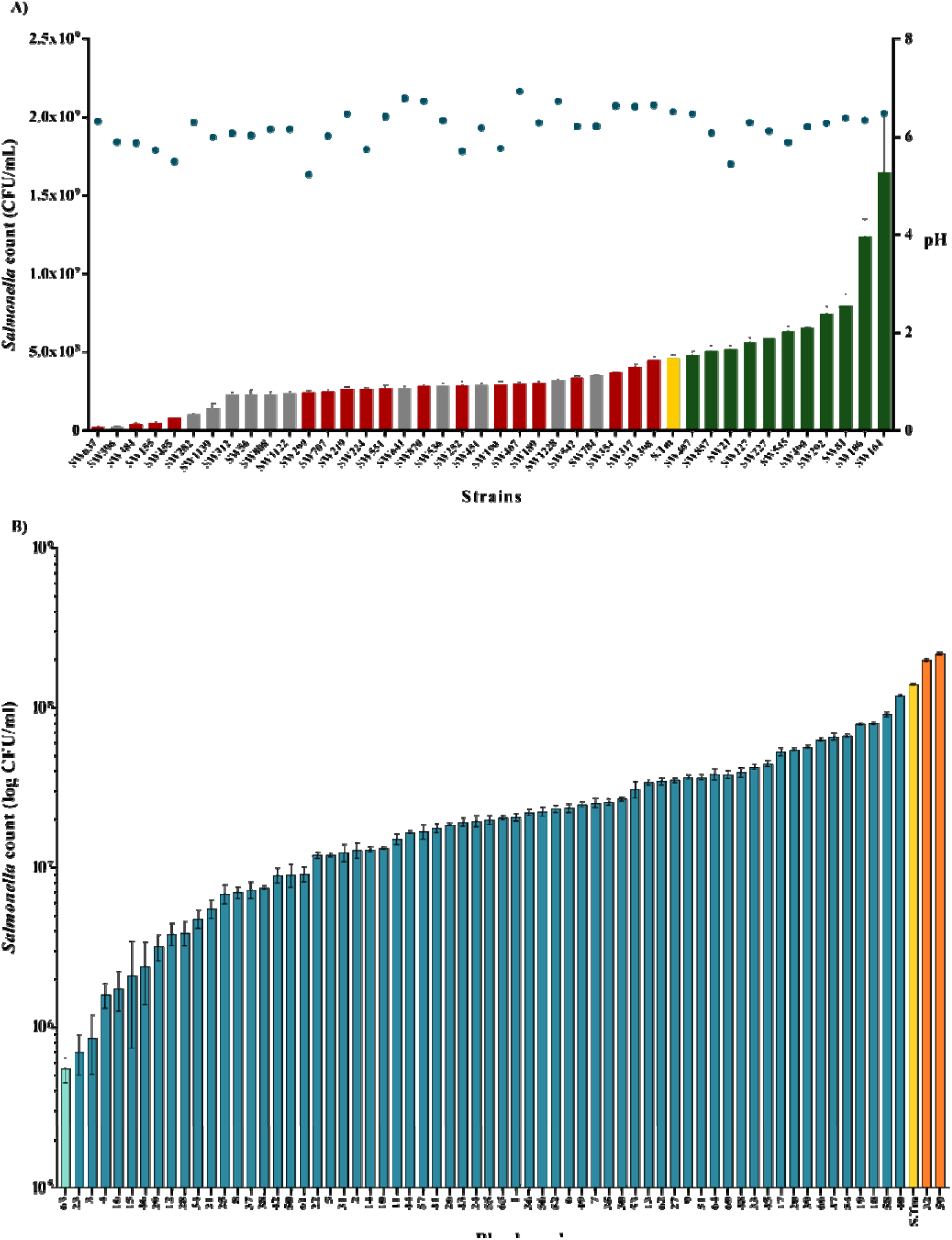
*In vitro* screening of representative species from the feral chicken gut microbiota library to identify *S*. Typhimurium inhibiting species. **A)** *Salmonella* inhibition capacity of individual species: Forty-one species isolated from the pool cecum of feral chickens were used for co-culture assays in this experiment. The OD_600_ of overnight bacterial culture was adjusted to 0.5 and individual strains were mixed with *S*. Typhimurium at a ratio of 9:1. The CFU of *Salmonella* (left y-axis) and pH (right y-axis) were determined after 24 hours incubation. The *S*. Typhimurium inhibiting strains are shown as red bars. The *S*. Typhimurium growth enhancing strains are shown as green bars. Twelve strains (grey color bars) were chosen to generate 66 combinations containing 10 species. **B)** *Salmonella* inhibition capacity of the bacterial blends: All 66 combinations were tested against *Salmonella* using the same co-culture assay described above. The *Salmonella* inhibiting blends are shown as blue bars. The *S*. Typhimurium growth enhancing blends were shown as orange bars. The blend showing highest inhibition level (light blue bar) is referred to as Mix10. All assays were performed in triplicate.

### Mix10 consortium confers partial protection against S. Typhimurium infection

We further determined the effect of Mix10 colonization on host health and *in vivo* inhibition capacity. To this end, we used a gnotobiotic chicken (*Gallus gallus*) model previously developed by our group [28] and a conventional chicken model. Briefly, we pooled an equal number of each species in Mix10 and used 10^7^ CFU/bird for inoculation while we used 10^5^ CFU/bird of *S*. Typhimurium for infection. Birds were euthanized at day 2 and day 5 post-infection (Fig. 3A). *Salmonella* load was determined from the cecum content. Overall, *S*. Typhimurium loads in each group trended to rise at day 2 and lower at day 5 post-infection, excepted Mix10-colonized gnotobiotic group that showed no difference of *S*. Typhimurium loads between day 2 and day 5 post-infection (Fig. 3B). At day 2 post-infection, *S*. Typhimurium loads in Mix10-colonized gnotobiotic group with *S*. Typhimurium infection were significantly lower than in the gnotobiotic group with *S*. Typhimurium infection (6.3-fold mean reduction). Although there was no significant different at day 5 post-infection, Mix10 reduced 1.4-fold of *S*. Typhimurium compared to the gnotobiotic group with *S*. Typhimurium infection. Mix10 also decreased *Salmonella* in conventional chickens, in which Mix10-colonized conventional group showed 2.4-fold of *S*. Typhimurium reduction compared to conventional group with *S*. Typhimurium infection at day 5 post-infection (Fig. 3B). This implicated that Mix10 support the resident bacteria to inhibit *Salmonella* colonization in chicken gut. Additionally, Mix10 also significantly lowered *S*. Typhimurium load in both gnotobiotic and conventional chickens compared to Mix59 which was observed to enhance growth of *S*. Typhimurium, corresponding to *in vitro* experiment (Fig. S2). The reduction of *Salmonella* load in Mix10 colonized groups was in line with our expectation that this consortium could inhibit *Salmonella in vivo*. Then, we examined the effect of Mix10 colonization on intestinal physiology via histopathology. Inflammatory symptoms of cecal tissues were evaluated using histological sections (Fig. 3C). Fibrinopurulent exudate was observed in the lumen of gnotobiotic group with *S*. Typhimurium infection (Fig. 3C; i). Also, the mucosa was swollen due to mixed inflammatory cell infiltrates such as macrophages, lymphocytes, and heterophils in lamina propria. Erosion of mucosa was evident with the loss of mucosal folds indicating granulomatous transmural enteritis. This deeper inflammation is typical of salmonellosis. Under higher magnification, early transmural inflammation with minimal peritonitis was observed in the Mix10-colonized gnotobiotic group with *S*. Typhimurium infection (Fig. 3C; ii). Inflammation of the mucosa was still detected which narrowed the luminal space, nonetheless, the severity of the infection was reduced in this group. Mucosal folds were noticeable but mixed inflammatory cells were still spotted. The mucosa was not eroded, and no exudate was found in the lumen. Mix10-colonized gnotobiotic chickens showed a large empty lumen with a small amount of ingesta (Fig. 3C; iii). Thin mucosa with mucosal folds was protruding into the lumen. Mild cellularity of lamina propria with scattered glands was observed. The intestinal epithelium of this group was improved towards that of conventional chicken (Fig. 3C; iv). In conventional chicken model, the highest inflammation symptoms were observed in conventional group with *S*. Typhimurium infection (Fig. 3C; v). Mix10-colonized conventional chicken with *S*. Typhimurium infection exhibited only a small amount of exudate in the lumen, and submucosal and transmural edema with macrophages and heterophils (Fig. 3C; v). When these histopathological images were compared, mucosal inflammation was very high in chicken with *S*. Typhimurium infection but subsided in groups with Mix10 colonization (Fig. 3D). *S*. Typhimurium infection in gnotobiotic and conventional group showed improved histopathology scores at day 5 post-infection, while Mix10 resulted in the least lesions as depicted by the histopathological scores compared to *S*. Typhimurium infection. The Mix10-colonized gnotobiotic group showed lower histopathological score compared to gnotobiotic group with *S*. Typhimurium infection that elevated score at day 5 post-infection. When Mix10 was administrated into gnotobiotic group, low score was observed at day 2 post-infection, and this score was reduced at day 5 post-infection. This suggested that Mix10 has capability to maintain gut physiology of gnotobiotic chicks. Mix10-colonized conventional group reduced histopathological scores compared to gnotobiotic group with *S*. Typhimurium infection. When histopathology results of Mix10 and Mix59 were compared to control, the inflammation of cecal tissue was lower in Mix10 administrating groups, whereas it was raised in Mix59 administrating groups, corresponding to the increase number of *Salmonella* in Mix59 in vitro and in vivo evidences (Fig. S4). These results suggested that Mix10 normalize chicken gut by supporting the development of intestinal tissue and reducing inflammatory symptoms and intact mucosa during *S*. Typhimurium infection.

**Figure 3.**
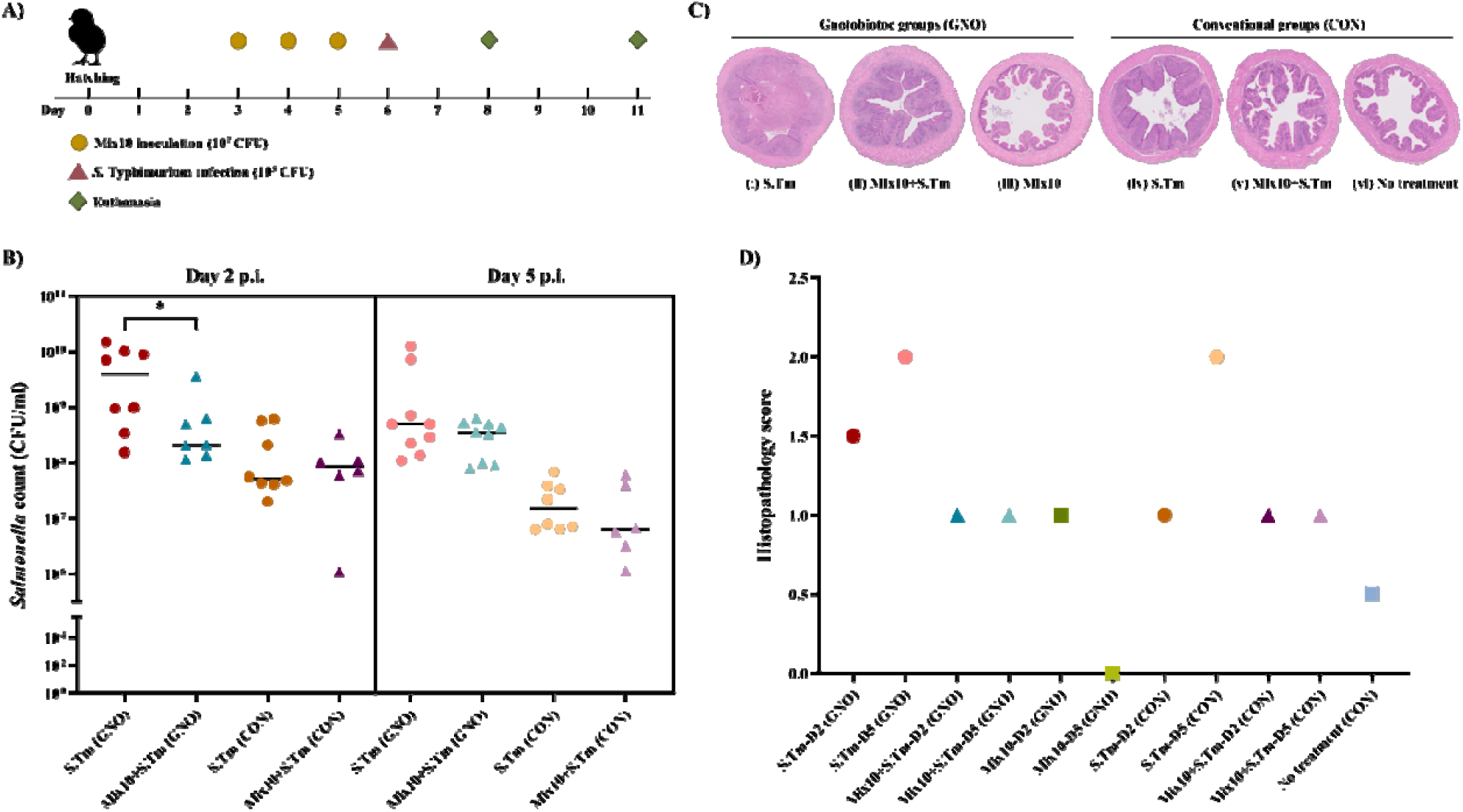
*In vivo* effect of Mix10 tested in a gnotobiotic chicken model. **A)** Chicken experimental design using Mix10 against *S*. Typhimurium *in vivo*: At hatching, the gnotobiotic chickens were divided into six different groups representing Mix10-colonized gnotobiotic group; Mix10 (GNO), Mix10-colonized gnotobiotic group with *S*. Typhimurium infection; Mix10+S.Tm (GNO), gnotobiotic group with *S*. Typhimurium infection; S.Tm (GNO), conventional group; no treatment (CON), Mix10-colonized conventional group with *S*. Typhimurium infection; Mix10+S.Tm (CON), and conventional group with *S*. Typhimurium infection; S.Tm (CON). Mix10 at 10^7^ CFU was administered via oral drenching at day 3, 4 and 5 post-hatching. Chickens were infected with 10^5^ CFU of *S*. Typhimurium. Half the number of chickens in each group were euthanized at day 2 post-infection, and others on day 5 post-infection. **B)** *S*. Typhimurium load in the infected groups: cecum content of gnotobiotic group with *S*. Typhimurium infection (n=17), Mix10-colonized gnotobiotic group with *S*. Typhimurium infection (n=16), conventional group with *S*. Typhimurium infection (n=16) and Mix10-colonized conventional group (n=12) with *S*. Typhimurium infection (n=10) on day 2 and 5 post-infection. Significant difference between control and Mix10-colonized groups was performed using the Mann-Whitney test; ^*^*P* < 0.05, ^**^*P* < 0.01, ^***^*P* < 0.001, and ^****^*P* < 0.0001. **C)** H&E stained histopathology of the cecum at day 11: (i) gnotobiotic group with *S*. Typhimurium infection; S.Tm (GNO), (ii) Mix10-colonized gnotobiotic group with *S*. Typhimurium infection; Mix10+S.Tm (GNO), (iii) Mix10-colonized gnotobiotic group; Mix10 (GNO), (iv) conventional group with *S*. Typhimurium infection; S.Tm (CON), (v) Mix10-colonized conventional group with *S*. Typhimurium infection; Mix10+S.Tm (CON), and (vi) conventional group; notreatment (CON) **D)** Histopathological scores: Cecal tissue samples of six groups at day 2 and day 5 post-infection were used to evaluate.

### Mix10 modulates gut immunity and reduces Salmonella-induced inflammation

In chicken, *Salmonella* infection is known to trigger inflammation of the gut by the production of pro-inflammatory cytokines such as interleukin (IL)-1 and IL-6 [29-32], chemokines such as IL-8, and type 1 T helper (Th1) cell cytokines such as IL-2 and interferon-*γ* (INF-*γ*), along with a cascade of other cytokines including tumor necrosis factor-*α* (TNF-*α*), IL-12, and IL-15 [33, 34]. As Mix10 was observed to inhibit *S*. Typhimurium *in vitro* and *in vivo* experiment, we investigated whether Mix10 colonization could ameliorate *Salmonella*-induced inflammation, the expression level of 84 inflammation-associated genes were measured by quantitative reverse-transcriptase polymerase chain reaction (qRT-PCR) array in the chicken caecum (Table S5). As expected, gnotobiotic chickens with *S*. Typhimurium infection showed multiple fold increase in the expression levels of various pro-inflammatory cytokines and chemokines; IL-18, IL-1*β*, IL-6 and IL-8L1 at day 5 post-infection. Massive expression of these pro-inflammatory cytokines were correlated to upregulated expression of other genes such as Toll-like receptors (TLRs), Nucleotide-binding oligomerization domain containg 1 (NOD1), Myeloid defferentiation primary response gene 88 (MyD88) which are cell surface pattern receptor recognitions (PPRs) and activators of inflammatory pathways suggesting capability of *S*. Typhimurium in the induction of inflammation in microbiota-free chickens. Conversely, *S*. Typhimurium infection in Mix10-conlonized conventional chickens exhibited downregulated expression of most genes particularly NFKB1, MAPKs and IRFs, mediators in the expression of inflammatory cytokines. These results demonstrated that microbiota-induced immunity maturation limit *Salmonella* induced inflammation. Furthermore, the chickens colonized with Mix10 did not generate a severe inflammatory response as the expression levels of pro-inflammatory cytokines (IL-6 and IL-18) was comparatively low. Antimicrobial peptides (AMPs) are crucial for eliminating a broad range of pathogens through pathogen-associated molecule pattern (PAMP) receptors. Two AMPs; cathelicidin2 (CATH2) as well as defensin-beta 1 (DEFB1) have been reported. In this study, Mix10-colonized chickens with *S*. Typhimurium infection showed a higher level of CATH2 as well as DEFB1 when compared to the group infected with *S*. Typhimurium (Fig. 4). The results suggested that colonization of Mix10 species in the chicken gut can ameliorate *S*. Typhimurium induced inflammation by activating AMPs production and anti-inflammatory immune response.

**Figure 4.**
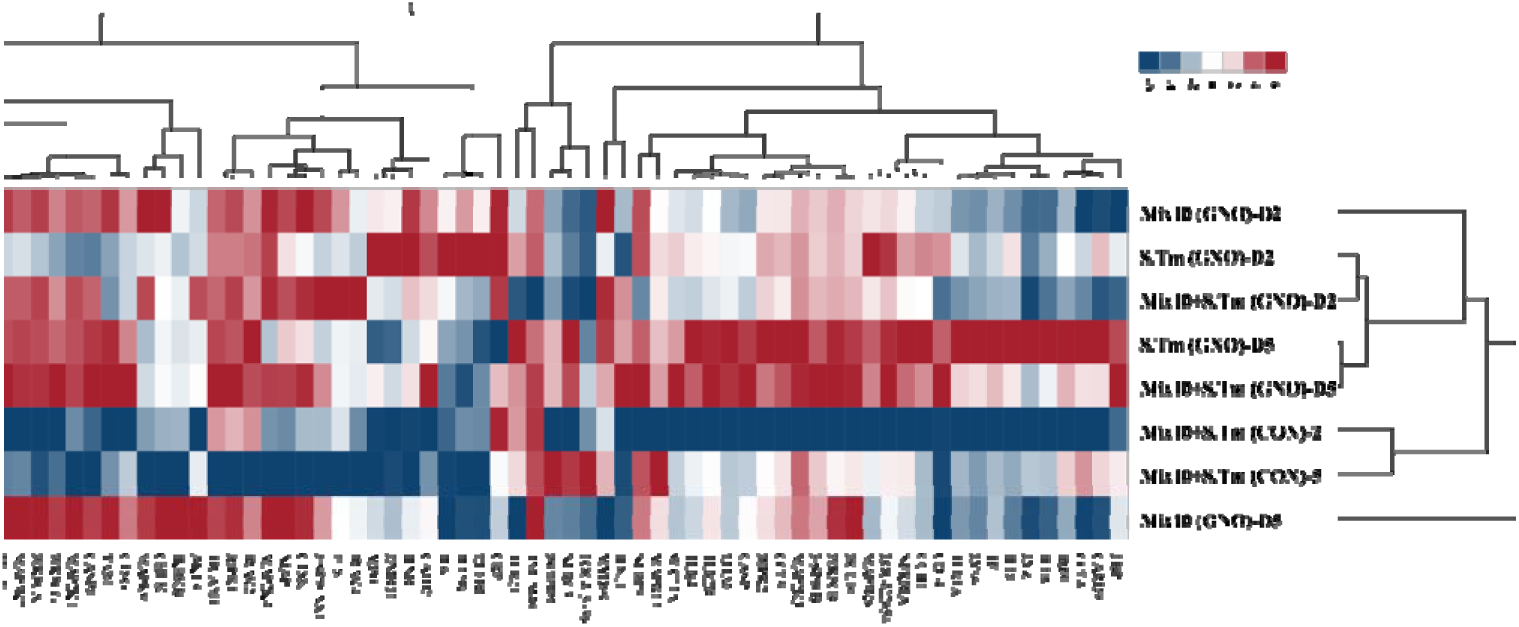
Determination of the immune response in chicken during Mix10 inoculation and *S*. Typhimurium infection. Total RNA from pooled cecal tissue from 4 groups: gnotobiotic group with *S*. Typhimurium infection; S.Tm (GNO), Mix10-colonized gnotobiotic group; Mix10 (GNO), Mix10-colonized gnotobiotic group with *S*. Typhimurium infection; Mix10+S.Tm (GNO), and Mix10-colonized conventional group with *S*. Typhimurium infection; Mix10+S.Tm (CON) was used to determine the inflammatory response. Data were presented as normalized fold change compared to gnotobiotic chicken as the baseline. Relative expression of the innate immune response, pro-inflammatory response and anti-inflammatory response at day 2 and 5 post-infection in all groups was compared.

### Mix10 in vivo community composition and functional genomic analysis

The microbial community profile of Mix10 colonization in chicken gut was determined using 16S rRNA amplicon from the cecal samples. Although all species of Mix10 was inoculated in equal proportion to gnotobiotic and conventional chickens, some species reached high abundance while others have low abundance or did not colonize the gut at all (Fig. 5 and Table S6). *Olsenella* sp., *Pseudoflavonifratctor* sp., and *Megamonas funiformis* together constituted more than 70% of Mix10 population in the cecum of Mix10-colonized chickens and Mix10-colonized chickens with *S*. Typhimurium infection. Conversely, *Staphylococcus saprophyticus, Bacillus paralicheniformis*, and *Enterococcus durans* were not detected in the samples suggesting that they did not successfully colonize chicken gut. The abundance of *Salmonella* in all Mix10-colonized groups compared to control groups was substantially lower (Fig. 5). This supported the reduction of *Salmonella* determined by CFU enumeration (Fig. 3B). However, differences in community composition were observed in Mix10-colonized conventional group compared to Mix10-colonized gnotobiotic group. The abundance of *Megamonas* was increased, while *Pseudoflavonifractor* was decreased. Furthermore, *Olsenella* whose abundance was very high in gnotobiotic group, was lost in conventional group.

**FIG 5.**
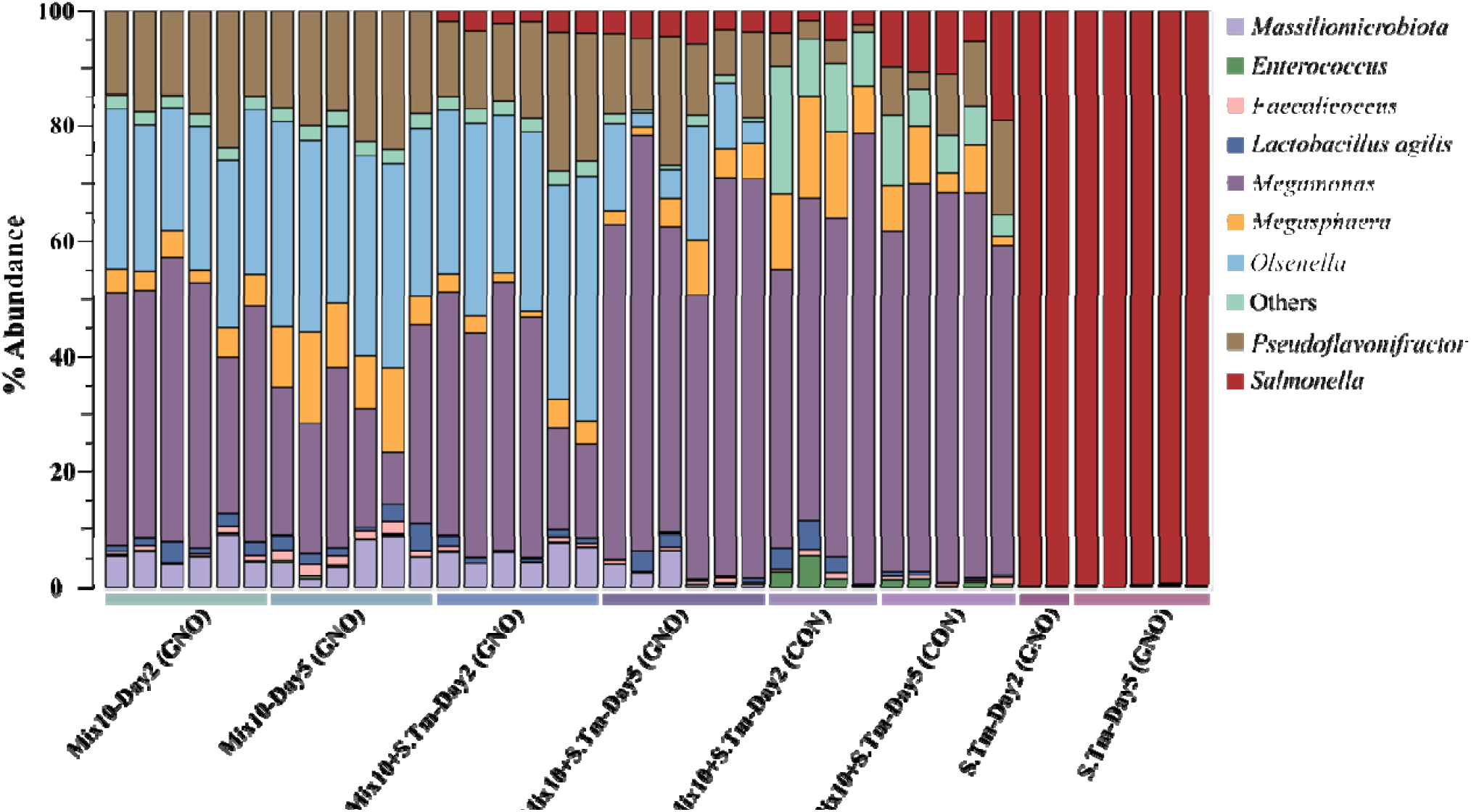
Mix10 population structure in chicken cecum determined using 16S rRNA sequencing. The relative abundance of individual species in the Mix10 after colonizing the gnotobiotic and conventional chicken and during *S*. Typhimurium infection were determined using 16S amplicon sequencing. The OTU clustering was performed at 97% similarity level using CLC Genomics Workbench (version 11.0.1) with the Greengenes database and a custom database of full length 16S rRNA gene sequences of Mix10 and *Salmonella*. The stacked bar plots of relative abundance at genus and species level (color code) were generated using Explicet software (version 2.10.5).

To decipher the overall functional capabilities of the members of Mix10, the genomes were sequenced and analyzed. The overall genomic attributes are presented in Table 1. Since the presence of functional modules computed using KEGG has been used to design defined gut bacterial belnds that partially inhibited *Salmonella* [35], we examined whether presence of KEGG modules correlated with *in vivo* colonization of strains in our study. The presence and completeness of KEGG modules in the strains were annotated and based on that a total of 293 KEGG modules were present either completely or partially across 10 species of Mix10 (Fig. 6A and Table S7). Based on the results from amplicon sequencing, only 7 organisms; *Olsenella, Pseudoflavonifratctor, Megamonas, Megasphaera, Massiliomicrobiota, Faecalicoccus* and *Lactobacillus* were able to colonize the gnotobiotic chicken gut with diverse abundance, in which they harbored a total of 234 modules including 122 complete modules. However, out of 293 modules detected across all strains, 243 modules with 159 complete modules were contributed by *Bacillus, Enterococcus* and *Staphylococcus* which did not colonize the gnotobiotic chicken gut. This indicates that presence of KEGG modules in the genome of Mix10 species may not be the primary determinant of colonization ability in the chicken gut. This was further evident when all functional modules in colonized and non-colonized strains of Mix10 were compared against the predicted complete modules in the feral chicken fecal metagenome (Fig. 6B). Although colonized strains clustered closer to metagenome of the feral chicken microbiome, presence or absence of KEGG module did not reveal any clear partitioning in this comparison.

**FIG 6.**
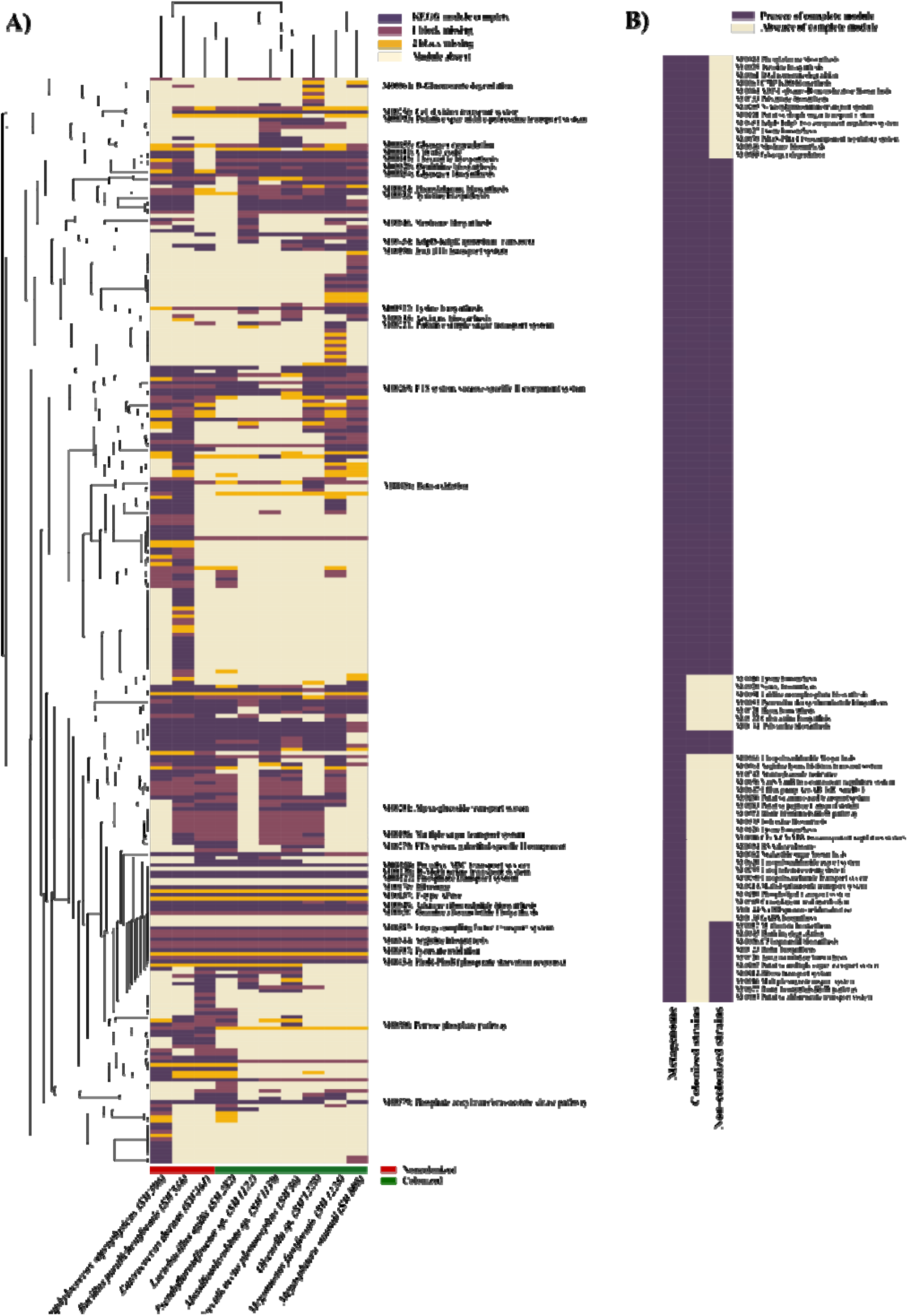
Functional clustering of KEGG modules present in Mix10 strains, and metagenome. To investigate the functional potential of all the strains, present in Mix10 and metagenome, the coding amino acid sequences were searched against the KO database against BlastKOALA and GhostKOALA server. **A)** The functional units of individual strains in Mix10 were presented by KEGG modules with 4 color scale: complete, 1 block missing, 2 blocks missing and module absent. **B)** All complete modules presented in metagenome data was compared to the set of colonized and non-colonized stains of Mix10. The matrix was then used to generate the heat maps using Pearson correlation and average linking method. The color code indicates the presence and completeness of each KEGG module. A few important KEGG module pathways were indicated in the heatmap. An extended list of KEGG modules and clusters are provided in supplementary table 7.

## Discussion

A mature microbial community in the adult gut is highly diverse and generally prevents the colonization of the pathogens such as *Salmonella*. The pathogen exclusion ability of the gut microbiota has been used to suppress *Clostridium difficile* infection in humans [7] and to exclude *Salmonella* in young chicken by fecal transplantation from healthy adults [8]. This capacity of the healthy microbiota to exclude or keep the pathogen numbers extremely low in the gut has been termed as competitive exclusion or colonization resistance [36]. The incredible complexity of the gut microbiota offers avenues to obtain and identify specific species or defined combination of species that can inhibit pathogens such as *Salmonella*. Recently, there have been propositions to do so using a modified Koch postulate[24, 25, 37]. As per this proposition, first the commensal organism is isolated as a pure culture, screened for pathogen exclusion capacity, and in the second stage, the ability of single species or defined communities of the isolated commensals to ameliorate disease is proven by mono or poly-associating them in a germfree host [38].

In this study, we used a similar approach to identify *Salmonella* inhibiting species from chicken gut microbiota. Since feral chicken has more microbial exposure than commercial chicken, we hypothesized that feral chicken microbiome could contain a high number of species that could provide colonization resistance against *Salmonella*. Using a culturomics based approach [39-42], 1,300 strains were isolated, and the species identity of the strains was determined. This collection was composed of 51 species (Fig. 1). From these, a co-culture assay-based screen identified 30 species which inhibited *Salmonella* in varying degrees (Fig. 2A). A modified Koch postulate to study microbiota function or mechanism proposes the concept of defined bacterial consortium as a unit that could ameliorate disease[12, 43] [24]. Simply put, this paradigm replaces “one pathogen = disease” with “one defined bacterial consortium = no disease”. We reasoned that the *Salmonella* inhibition ability of individual species could be improved if a defined consortium is made by pooling several strains. Consistent with this expectation, a pool of 10 species showed several fold higher *Salmonella* inhibition capacity (Fig. 2B). What we also observed here is that two out of the 66 blends increased the *Salmonella* growth *in vitro* and *in vivo* (Fig. 2B, Fig. S3 and Fig. S4). This is a clear demonstration of the fact that bacterial interaction can change individual strain phenotype, for instant *Salmonella* inhibition and is consistent with the concept of “defined bacterial consortium” as a unit to interrogate microbiota function as proposed in the modified Koch postulate for micorbiota [24, 25].

The blend (Mix10) that showed maximum *Salmonella* inhibition was then tested in a gnotobiotic and conventional chicken model [28] to determine whether it could exclude *Salmonella* in the chicken gut and reduce *Salmonella*-induced disease. Our results clearly showed that Mix10 partially excluded *Salmonella* (Fig. 3B), reduced tissue damage (Fig. 3C and D), and substantially reduced inflammation (Fig. 4,) in the chicken gut. These results were the validation of the second component in the modified Koch postulate, *i*.*e*., a demonstration that defined microbial consortium can ameliorate pathogen induced disease, such as *Salmonella*. Our results also showed that this inhibition is not serotype dependent and is applicable to other serotypes of *Salmonella* that commonly colonize chicken (Fig. S1). Moreover, the cell free supernatant of the Mix10 failed to inhibit *Salmonella* (Fig S2). On the basis of these results, the mechanism of Mix10 against *Salmonella* colonization could be nutrient competition of species or improvement of host immune system. When the Mix10 population structure was determined using 16S rRNA amplicon sequencing, we found that out of ten inoculated species, three did not colonize (*Staphylococcus saprophyticus, Bacillus paralicheniformis*, and *Enterococcus durans*) while three (*Olsenella* sp., *Pseudoflavonifratctor* sp., and *Megamonas funiformis)* dominated the population constituting more than 70% abundance in the cecum. Since these three species dominated all groups, we asserted that they are the key members of Mix10 consortium which produce most of the disease reduction in gnotobiotic chicken. However, loss of colonization may be because of host selection via adhesive molecule such as mucus glycans and immunoglobulin A, and immune system such as antimicrobial compounds; RegIII*γ* and defensis [44-47]. Overall, our approach of making defined bacterial consortium from a pure culture library and testing them in a native germfree host offers a means to select a subset of strains for further enhancement by excluding non-colonizers and selecting dominating species in the tested consortium.

Eventhough Mix10 may not be the best possible consortium of defined bacteria that could exclude *Salmonella* in chicken, it is possible that many such defined consortia of same or better effectiveness could be formulated from a pure culture library of isolates. As per the redundancy or insurance hypothesis, more than one species is retained in the gut ecosystem to ensure that loss of one species does not result in the loss of function contributed to microbiome by that species [3, 48]. Therefore, it is more than likely that many species exist in chicken microbiota contribute to the same function and their inhibition property can change depending on the composition of the consortium. When the dominating species from several blends such as Mix10 are pooled or stacked, better combinations may emerge. Nevertheless, our results show that choosing a diverse microbiota source is essential to gain *Salmonella* inhibiting species because two of three highly dominating species in Mix10 were previously uncultured species, justifying our choice of feral chicken as the input for culture library development. The previous study showed that the use of microbial diversity in the gut as a guide to developing *Salmonella* inhibiting defined consortium that was tested in a gnotobiotic mouse model [35]. However, we did not find a good correlation between the presence of KEGG modules in the genome and colonization of the strains *in vivo*. In the present study, we have tested only representative species in our library. Since other strains in the same species may have the inhibitory capacity, formulating additional blends from such strains in our library, performing combinatorial *in vitro* inhibition asasay followed by testing in the gnotobiotic chicken model, and pooling highly dominating species from multiple mixes may help to design well defined microbial ecosystem therapeutic against *Salmonella*.

## Material and Methods

### Development of the feral chicken gut microbiota library

Protocols used in this study for sample collection and chicken experiments were reviewed and approved by the Institutional Animal Care and Use Committee (IACUC) at the South Dakota State University, Brookings, South Dakota. For the isolation of bacteria from the feral chicken gut, six intestinal samples were pooled. The pooled intestinal sample was serially diluted and was plated on modified Brain Heart Infusion agar (BHI-M) with 12 different selective conditions (Supplementary Table 1). The modified Brain Heart Infusion agar (BHI-M) contained the following ingredients: 37 g/L of BHI, 5 g/L of yeast extract, 1mL of 1 mg/mL menadione, 0.3 g of L-cysteine, 1 mL of 0.25 mg/L of resazurin, 1 mL of 0.5mg/mL hemin, 10 ml of vitamin and mineral mixture, 1.7 ml of 30 mM acetic acid, 2 ml of 8 mM propionic acid, 2 ml of 4 mM butyric acid, 100 µl of 1 mM isovaleric acid, and 1% of pectin and inulin. All cultures were performed inside an anaerobic chamber (Coy Laboratories) containing 85% CO_2_, 10% H_2_, and 5% N_2_ maintained at 37^°^C. Total of 1,300 colonies was picked from all conditions and dilutions based on colony morphologies. Selected colonies were streaked on base BHI-M agar, and a single colony was selected for preparing stocks and species identification. Species identity of the isolates was determined using Matrix-Assisted Laser Desorption/ Ionization-Time of Flight (MALDI-TOF) or 16S rRNA gene sequencing. For MALDI-TOF identification, a single colony was smeared on the MALDI-TOF target plate and lysed by 70% formic acid. MALDI-TOF targets were covered with 1 µL of a matrix solution. MALDI-TOF was performed through Microflex LT system (Bruker Daltonics). A MALDI-TOF score >1.9 was considered as positive species identification. Isolates that could not be identified at this cut-off were identified using 16S rRNA gene sequencing. To identify these isolates, genomic DNA of overnight culture from a single colony was extracted using a DNeasy Blood & Tissue kit (Qiagen), according to the manufacturer’s instructions. Then 16S rRNA gene sequences were amplified using universal primer set 27F (5’-AGAGTTTGATCMTGGCTCAG-3’; Lane et al., 1991) and 1492R (5’-ACCTTGTTACGACTT-3’; Stackebrandt et al., 1993) [49, 50], and sequenced using a Sanger DNA sequencer (ABI 3730XL; Applied Biosystems) using 27F primer. The 16S rRNA gene sequence was used to verify species using the GenBank (www.ncbi.nlm.nih.gov/genbank/) and EZBioClound databases (www.ezbiocloud.net/eztaxon) [26]. All identified isolates were maintained in BHI-M medium with 10% (v/v) Dimethyl Sulfoxide (DMSO) at −80^°^C. Aerotolerance of the bacterial species was tested by culturing in aerobic, anaerobic and microaerophilic conditions. To this end, individual bacteria were first cultured overnight in BHI-M broth at 37^°^C under anaerobic condition. The optical density at 600 nm (OD_600_) of the cultures was adjusted to 0.5. Then, 1% of OD_600_ adjusted cultures were inoculated in fresh BHI-M media in triplicates. Each replicate of cultures was then incubated under anaerobic, microaerophilic and aerobic conditions. For microaerophilic condition, a hypoxic box was used to incubate the culture. After 24 hours of incubation, the growth of individual bacteria was determined by measuring OD_600._

### Co-culture assays, formulation of the bacterial blends and determination of inhibitory mechanism

A co-culture assay was used to screen all bacterial species for *S*. Typhimurium inhibition capacity. The growth of each bacterial species was measured following overnight incubation in BHI-M using a spectrophotometer at OD_600_. Subsequently, bacterial cells were maintained by adjusting the OD_600_ to 0.5 with 10% (v/v) DMSO at −80^°^C. In this assay, each bacterial stock was anaerobically cultured together with *S. Typhimurium* in a ratio of 9:1 in 1 mL of BHI-M broth and incubated at 37^°^C for 24h. To quantify the magnitude of *S*. Typhimurium inhibition by each species, the individual co-cultures were 10-fold serially diluted with 1X anaerobic phosphate buffer saline (PBS) and plated on Xylose Lysine Tergitol 4 (XLT4) agar (BD Difco, Houston, TX). The plates were incubated aerobically at 37^°^C for 24 hours followed by plating on XLT4 agar and colony forming units (CFU) were enumerated to determine the degree of *S*. Typhimurium inhibition.

To test whether Mix10 could inhibit multiple serotypes of *Salmonella*, we determined the inhibitory capacity of Mix10 against four serovars *Salmonella* (*S*. Typhimurium, *S*. Heidelberg, *S*. Infantis and *S*. Enteritidis) that are frequently infect poultry. A co-culture assay was performed as previously described in the methods.

To determine whether Mix10 inhibits *Salmonella* through nutrient competition or through other mechanisms, cell-free supernatants were tested for *Salmonella* inhibition. To this end, Mix10 was grown cultured as before for 48 hours. The cell pellets were removed by centrifugation at 3,000 rpm for 1 h. The supernatant was then filtered through 0.4 *μ*m filter. The purified supernatant was adjusted pH to 6.5-6.8 using NaOH and HCl. The supernatant was divided into three fractions. The first fraction was untreated, the second fraction was heated at 100^°^C for one hour while the third fraction was treated with 50 *μ*g/mL of proteinase K for one hour at 37^°^C. Following this, *S*. Typhimurium was cultured in the three fractions diluted 50% with BHI-M. In parallel, equal dilution of BHI-M with 1X PBS was used to culture *Salmonella* as a control sample. After 24 hours of incubation, CFU of *Salmonella* was enumerated as previously described.

### Determination of in vivo effect of ten species consortium using gnotobiotic and conventional chicken model

We used a gnotobiotic chicken model developed by our group to determine the *in vivo* effect of Mix10. Briefly, the gnotobiotic chicks were hatched by following the previously described protocol [28]. Fertile Specific Pathogen Free (SPF) eggs were wiped with Sporicidin^®^ disinfectant solution (Contec^®^, Inc.), an FDA approved sterilizing solution, followed by washing in sterile water. Further, the eggs were incubated at 37^°^C and 55% humidity for 19 days. Eggs containing an embryo, confirmed after candling, were dipped in Sporicidin^®^ for 15s and wiped with sterile water before transferring to a biosafety cabinet maintained at 37^°^C and 65% humidity until hatching. Chickens were orally drenched with 10^7^ CFU of Mix10 (best Salmonella inhbiting mix) / Mix59 (Mix enhancing Salmonella growth) (Fig 2) at day 3, 4, and 5 post-hatching, followed by 10^5^ CFU of *S*. Typhimurium challenge on day 6 post-hatching (Fig 3A). Similarly, for the conventional group, eggs were not sterilized and were allowed to hatch normally. We fed chickens with 10^5^ CFU of *S*. Typhimurium and 10^7^ CFU of Mix10/Mix59 on day 6 post-hatching. Chickens were euthanized by cervical dislocation on day 2 and day 5 post-infection **(Fig. 3A)**. The cecum contents and tissues were aseptically collected for further analysis. *S*. Typhimurium load in the cecum contents were determined by plating on *Salmonella* selective XLT4 agar.

### Histopathology

The tissues for histopathology were initially fixed in 10% formalin. The cecum tissues were trimmed and processed into paraffin blocks by routine histopathological methods, *i*.*e*., gradual dehydration through a series of ethanol immersion, followed by xylene and then paraffin wax. They were sectioned at 4*μ*m and stained with hematoxylin and eosin (HE), followed by scanning of glass slides in a motic scanner. Further, the cecum pathology was evaluated based on scores.

### Assessment of immune response using Quantitative Reverse-Transcriptase PCR (Q-PCR)

To study the chicken immune response after bacterial treatment, gene expression in the cecal tissue was determined. Total RNA from cecal tissue samples was extracted using the TRIzol^®^ reagent (Ambion | RNA, Invitrogen) method. Briefly, an average weight of 0.042 g of cecal tissue per sample (n=7 per group) was used. Tissue samples from each group were pooled and homogenized separately in TRIzol^®^ reagent (1mL per 100 mg of tissue sample). RNA extraction was performed according to manufacturer’s protocol. RNA concentration was determined using spectrophotometric optical density measurement (A260/A280) by NanoDrop™ One (Thermo Fisher Scientific, Wilmington, DE). For Q-PCR, cDNA was synthesized using First-strand cDNA synthesis kit (New England BioLabs, Inc.) according to the manufacturer’s protocol. To get enough cDNA for downstream procedures, 4 µg of RNA was used as input in a cDNA synthesis. The dynamics of the chicken immune response was analyzed using RT2 Profiler PCR Array (cat# 8ZA-1214, Qiagen) according to the manufacturer’s protocol. Real-time q-PCR was performed following the manufacturer’s protocol using an ABI 7500HT thermal cycler (Applied Biosystems). A cycle threshold cut-off of 0.2 was applied to all gene amplifications and was normalized to Ribosomal protein L4 (RPL4) and Hydroxymethylbilane synthase (HMBS) as they were stably expressed across all treatment groups from a panel of five housekeeping genes. The fold regulation of genes in 4 treatment groups was calculated by comparing to control group (gnotobiotic chickens). Data was clustered using Pearson correlation with complete linkage in Morpheus package.

### Determination of the population structure of the bacterial consortium in the cecum using 16S rRNA amplicon analysis

We determined the relative abundance of individual species in the Mix10 after colonizing the gnotobiotic and conventional chicken using 16S rRNA amplicon sequencing. Genomic DNA from cecal contents was extracted using the PowerSoil DNA isolation kit (Mo Bio Laboratories Inc, CA). To ensure even lysis of the microbial community, bead beating was performed on 100 mg of cecal contents for 10 min using a tissue lyser (Qiagen, Germantown, MD). Remaining steps for DNA isolation were performed as per manufacturer’s instruction. Final elution of DNA was carried out in 50 µL of nuclease-free water. The quality of DNA was assessed using a NanoDrop^™^ One and quantified using a Qubit Fluorometer 3.0 (Invitrogen, Carlsbad, CA). The samples were stored at −20^°^C until further use. The samples carrying low DNA yield were removed from the downstream processes. The enrichment of the microbial DNA was performed using the NEBNext^®^ Microbiome DNA Enrichment Kit (New England Biolabs Inc, MA) according to the manufacturer’s instruction. A total of 31 DNA samples were used for 16S rRNA gene sequencing using the Illumina MiSeq platform with 250 base paired-end V2 chemistry. DNA library preparation was performed using Illumina Nextera XT library preparation kit (Illumina Inc. San Diego, CA) targeting the V3 and V4 region of the 16S rRNA gene sequence (Klindworth et al., 2013). The amplicons were then purified using Agencourt AMPure XP beads (Beckman Coulter). Before loading, libraries were bead normalized and pooled in equal concentration. After sequencing, CLC Genomics Workbench (version 11.0.1) (Qiagen) was used to analyze the 16S rRNA sequence data. An average of 72,749 raw reads per sample (ranging from 34,962 to 100,936) was imported to CLC workbench. After the initial quality check, reads with low Q30 score was removed by trimming with a quality score limit of 0.01. Paired reads were merged at a minimum alignment score of 40. OTU clustering was performed at the 97% similarity level using a locally downloaded Greengenes database [51] and a custom database of full-length 16S rRNA gene sequence of Mix10 species and *Salmonella*. Best matches were found at chimera cross over cost of 3 and kmer size of 6. Finally, on an average 28,759 reads per sample were used to generate OTUs. The abundance table and metadata were then used to create stacked bar plots in Explicet software tool (version 2.10.5) [52]. The plot was generated using only those OTUs (genus level) that have more than 0.25 percent relative abundance across all samples.

### Genome analysis of Mix10 species and metagenomic analysis of intestinal content using next-generation sequencing

We used the Bacterial DNA kit (D3350-02, e.Z.N.A™, OMEGA bio-tek, USA) to isolate the genomic DNA of individual species. The quality of DNA was assessed using Qubit Fluorometer 3.0. Whole genome sequencing was performed using Illumina MiSeq platform using MiSeq Reagent Kit V3 chemistry. The reads were assembled using Unicycler that builds an initial assembly graph from short reads using the de novo assembler SPAdes 3.11.1 [53]. The quality assessment for the assemblies was performed using QUAST [54]. The open reading frames (ORFs) were predicted using Prodigal 2.6 [55] in the Prokka software package [56]. To determine the functional modules in the genome, the amino acid sequences were mapped against the KEGG (Kyoto Encyclopedia of Genes and Genomes) database using the BlastKOALA genome annotation tool [57]. Each KEGG module was represented on a scale of 0 to 4 (0= complete, 1=1 block missing, 2= 2 block missing and 3= module absent). The matrix was used for hierarchical clustering using the Morpheus (https://software.broadinstitute.org/morpheus) server for constructing the heat map using Pearson correlation matrix and average linkage method. As mentioned previously, the strains of culture library were isolated from the pooled intestinal content of six feral chickens. This original sample was used for DNA isolation, sequencing and analysis for our previous study [28]. In this study, the assembled contigs from this inoculum were used to predict the putative protein coding sequences using FragGeneScan [58]. The resulting amino acid sequences were clustered using CD-HIT to reduce the sequence redundancy [59]. The clustered proteins were then annotated against the KEGG Orthology (KO) database to assign the molecular functions using GhostKOALA (PMID: 26585406). The complete modules present in the metagenomics sample were compared against the colonized (n=7) and non-colonized strains (n=3).

### Statistical analysis

Statistical analysis was performed using the exact Mann-Whitney (MW) U test using GraphPad Prism version 9.0.0 for Windows (GraphPad Software, www.graphpad.com). *P* values less than 0.05 were considered as statistically significant (^*^ *P* < 0.05, ^**^ *P* < 0.01, ^***^ P < 0.001, ^****^ *P*, 0.0001)

### Data availability

Draft genome of individual Mix10 strains and raw data of 16S rRNA amplicon metagenomics in this study were deposited in the NCBI under BioProject number PRJNA524186.

## Acknowledgements

Computations supporting this project were performed on High-Performance Computing systems managed by Research Computing Group, part of the Division of Technology and Security at South Dakota State University. This work was in part supported by the grants from the South Dakota Governors Office of Economic Development (SD-GOED) and the United States Department of Agriculture (grant numbers grant numbers SD00R646-18 and SD00H702-20) awarded to JS. SW received support from the Science Achievement Scholarship of Thailand.

## Supplementary Material

**Table S1. Media composition and supplements used for bacterial isolation**. BHI-M was used as a base medium in this study. Total 12 culture conditions; base BHI-M medium and 11 selective conditions by supplementing BHI-M with antibiotics were used to isolate bacterial strains. The concentration of antibiotics used in this study was selected from Minimum inhibitory concentration (MIC) data from the EUCAST.

**Table S2. List of bacterial strains isolated in this study**. Total 1,300 isolates were selected from the cultures of pooled intestinal content in 12 culture conditions. Species identity of 1,023 isolates was determined using MALDI-TOF or 16S rRNA gene sequencing.

**Table S3. Bacterial strains used in the individual co-culture assay against *S*. Typhimurium**. After species identification, 41 species representative isolates were co-cultured with *S*. Typhimurium in anaerobic condition. After 24 hours of co-culture, enumeration of *S*. Typhimurium was performed on *Salmonella* selective plate XLT4 agar, and pH of individual co-cultures was measured.

**Table** S**4. List of bacterial species used to generate defined blends and composition of the blends**. Twelve *S*. Typhimurium inhibiting, and fast-growing species were selected from the individual co-culture assay. They were used to generate combination by randomly removing 2 species at a time.

**Table** S5. **Immune response of gnotobiotic chicken during Mix10 colonization and *S*. Typhimurium infection**. Antibacterial immune response in the cecal tissue measured using qRT-PCR array at day 2 and 5 post infection. The normalization and analysis of the data were performed using GeneGlobe Data Analysis Center Web server.

**Table** S6. **OTUs clustering at genus level of Mix10 colonization in gnotobiotic chicken gut**. The table shows OTU at genus level present in 3 groups at day 2 and day 5 post-infection by CLC Genomics Workbench (version 11.0.1) (Qiagen). The color codes of columns correspond to groups in Fig. 5. The rows with yellow highlight referred to as genus shown in Fig. 5.

**Table** S**7. Functional clustering of KEGG modules of individual strains of Mix10**. The amino acid sequences were mapped against the KEGG (Kyoto Encyclopedia of Genes and Genomes) database using the BlastKOALA and GhostKOALA genome annotation tool. Each KEGG module was represented using the following scale: 0= Complete module, 1= 1 block missing, 2= 2 block missing and 3= module absent.

**FIG S1.**
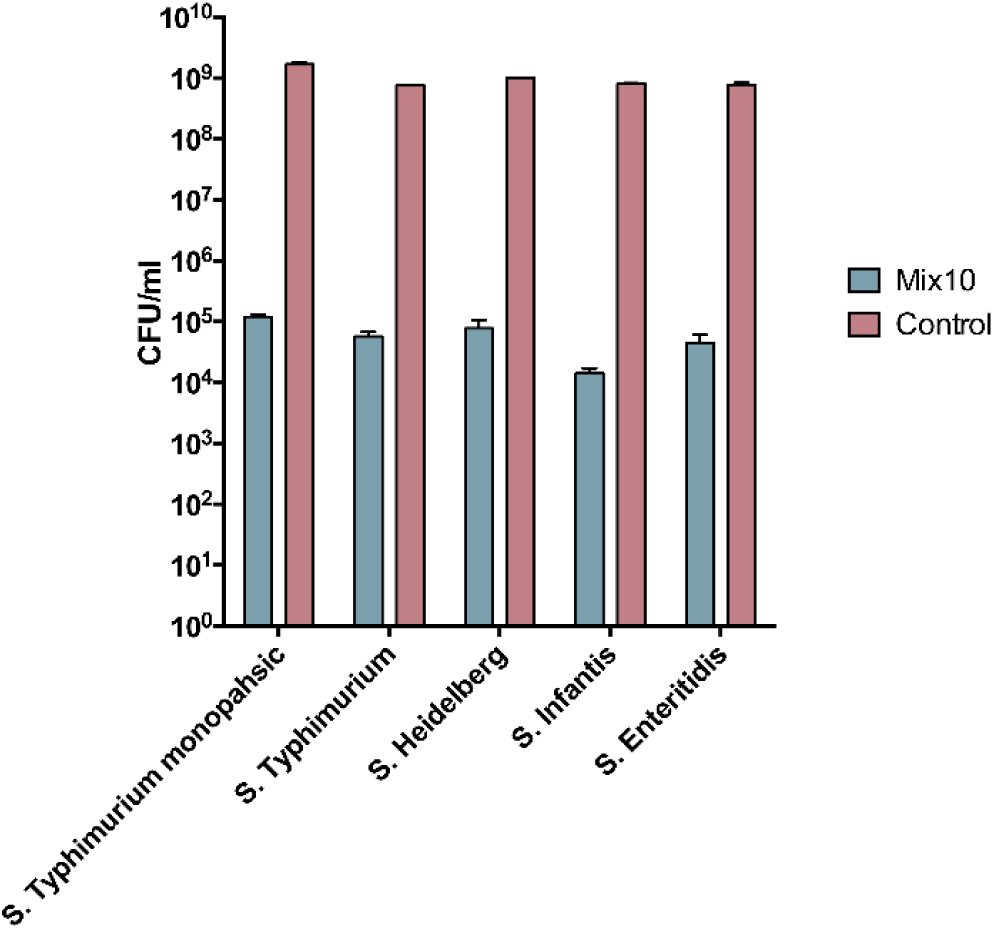
Mix10 inhibition of Salmonella serovars frequently found in poultry. Inhibitory capacity of Mix10 against serovars *S*. Typhimurium, *S*. Heidelberg, *S*. Infantis and *S*. Enteritidis were determined using the co-culture assay. The bars show CFU/ml of serotypes of *Salmonella* after 24 h of co-culture with Mix10 (blue bars) and *Salmonella* monoculture (pink bars).

**FIG S2.**
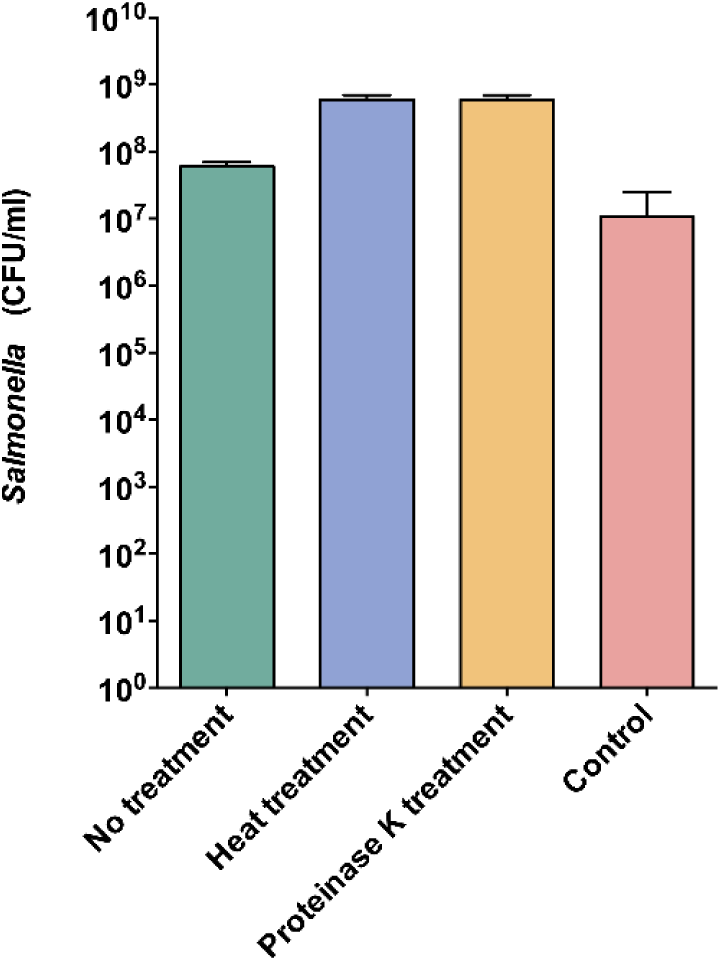
Inhibitory effect of Mix10 cell-free supernatant on *S*. Typhimurium. To determine the inhibitory mechanism of Mix10 against Salmonella, a cell-free supernatant co-culture assay was performed. Cell free supernant of Mix10 was prepared by centrifugation of Mix10 culture at 3,000 rpm for 1 hour. The supernatant was then filtered through 0.4 *μ*m filter. The purified supernatant was adjusted pH to 6.5-6.8 using NaOH and HCl. The supernatant was divided into 3 fractions; no treatment, heat treatment heated at 100^°^C for 1 hour, and 50 *μ*g/ml of proteinase K for 1 hour at 37^°^C. After 24 hours of incubation with cell free supernatant, *Salmonella* CFU was enumerated as described previously.

**FIG S3.**
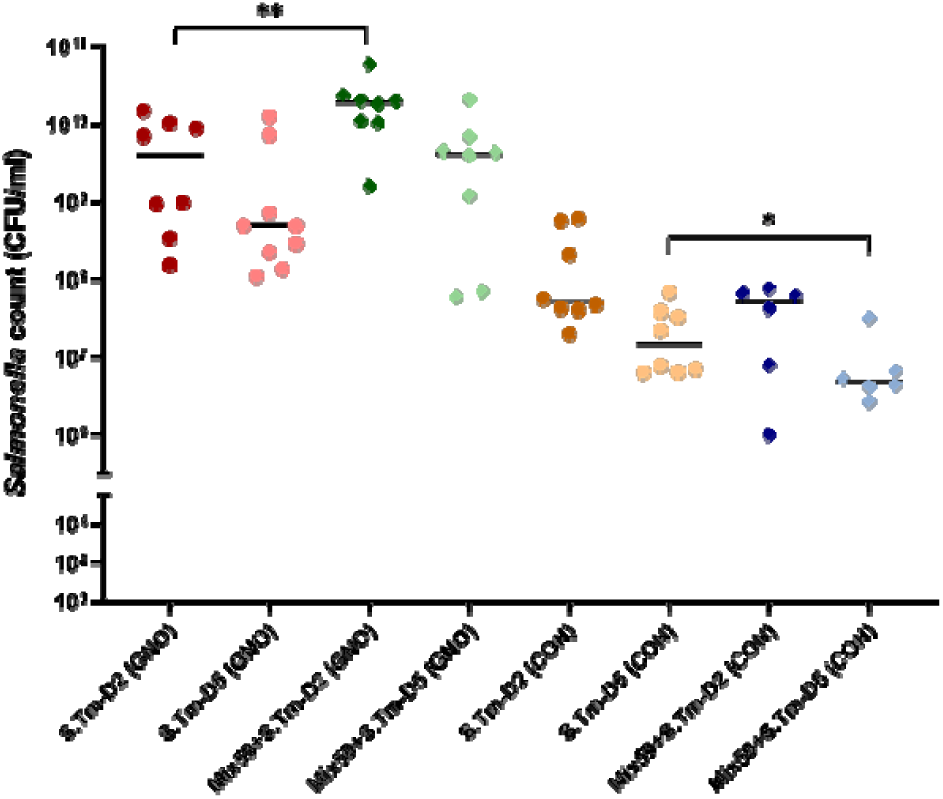
*Salmonella* load in *Salmonella* growth enhancing blend (Mix59) colonized chickens. Mix59 was designed to test against *S*. Typhimurium *in vivo*. The experiment platform was similar to the *in vivo* test of Mix10 (Fig. 3A). *S*. Typhimurium load in the infected groups: cecum content of gnotobiotic group with *S*. Typhimurium infection (n=17), Mix59-colonized gnotobiotic group with *S*. Typhimurium infection (n=16), conventional group with *S*. Typhimurium infection (n=16) and Mix59-colonized conventional group with *S*. Typhimurium infection (n=12) on day 2 and 5 post-infection. Significant difference between control and Mix59-colonied groups was performed using the Mann-Whitney test; ^*^*P* < 0.05, ^**^*P* < 0.01, ^***^*P* < 0.001, and ^****^*P* < 0.0001.

**FIG S4.**
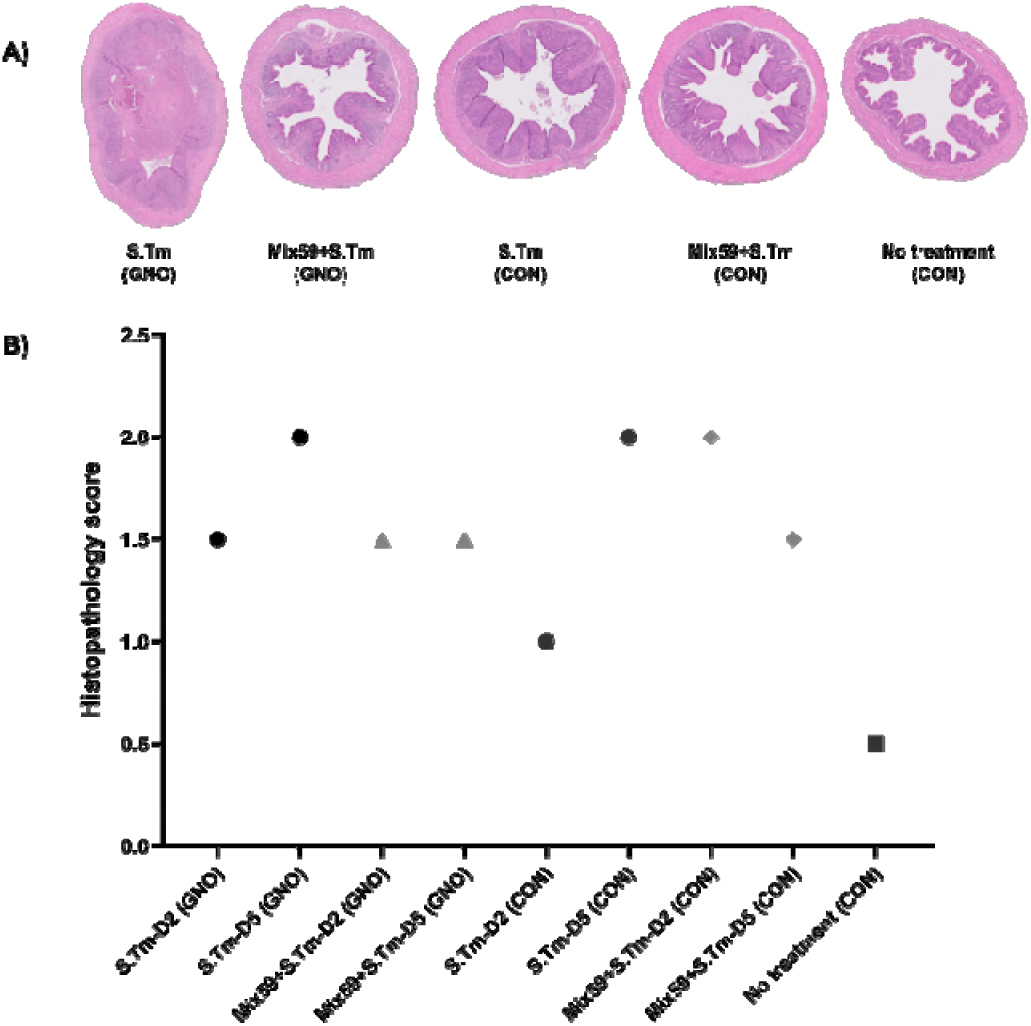
Histopathology of the cecum. **A)** Cecum of gnotobiotc and conventional chickens were performed H&E staining; gnotobiotic group with *S*. Typhimurium infection; S.Tm (GNO), Mix59-colonized gnotobiotic group with *S*. Typhimurium infection; Mix59+S.Tm (GNO), conventional group with *S*. Typhimurium infection; S.Tm (CON), Mix59-colonized conventional group with *S*. Typhimurium infection; Mix59+S.Tm (CON), and conventional group; notreatment (CON) **B)** Histopathological scores: Cecal tissue samples of 5 groups were used to evaluate.

